# Motor-mediated clustering at microtubule plus ends facilitates protein transfer to a bio-mimetic cortex

**DOI:** 10.1101/736728

**Authors:** Núria Taberner, Marileen Dogterom

**Affiliations:** Department of Bionanoscience, Kavli Institute of Nanoscience, Delft University of Technology, Delft, The Netherlands; RIKEN BDR, Kobe, Japan

## Abstract

Polarized protein distributions at the cortex play an important role in the spatial organization of cells. In *S. pombe*, growing microtubule ends contribute to the establishment and maintenance of such distributions by delivering specific factors to membrane receptors at the poles of the cell. It is however unclear how microtubule plus-end tracking of proteins favours protein accumulation at the cell cortex compared to proteins arriving directly from the cytoplasm. To address this question, we developed an *in vitro* assay, where microtubules were made to deliver His-tagged plus-end tracking proteins to functionalized microchamber walls. We found that motor-mediated protein clusters formed at microtubule ends were able to transfer to the walls, but non-clustered proteins were not. We further show that this transfer mechanism leads to preferential cluster accumulation at chamber poles, when microtubules are confined to elongated microfabricated chambers with sizes and shapes similar to *S. pombe*.

## Introduction

Polarized distributions of proteins at the cell cortex serve as cues to organize and localize intracellular components. Such protein distributions are involved in defining cell polarity and dictating the directionality of global processes such as cell division, migration, or polarized cell growth [Nelson 2003, Macara and Milli 2010]. There are multiple mechanisms by which cortical protein distributions may in principle be formed, either based on reaction-diffusion principles and/or dependent on cytoskeletal processes [Vendel et al. 2019], but the exact mechanisms are not yet understood. Here we focus on the principles behind establishment of polarized protein distributions in the model system fission yeast, where dynamic microtubules are essential in targeting cell-end specific factors to the cell poles.

Microtubules are long protein filaments formed by the assembly of alpha and beta-tubulin heterodimers. They typically consist of 13 protofilaments that form a hollow tube of about 24 nm diameter that can extend up to several tens of micrometres [Tilney et al. 1973]. Microtubules stochastically switch between two states in a process termed dynamic instability: a slow-growing state caused by the net addition and lateral association of heterodimers at the protofilament ends; and a fast-shrinking state caused by curling out of the protofilament ends and subsequent disassembly from the tubular structure into short unstable pieces [Mitchison and Kirschner 1984, Desai & Mitchison 1997, Walker et al. 1988]. The switch from growth to shrinkage is termed a ‘catastrophe’ and the reverse a ‘rescue’. The end of the microtubule that is exposing beta-tubulins, termed the plus end, is highly dynamic. The other end, the minus end, is much less dynamic and is often anchored at a microtubule nucleating centre.

In cells, microtubules typically grow from nucleating centres near the middle of the cell towards the cell periphery, where contact with the cell cortex may cause temporal stalling or buckling, followed by catastrophe [Tran et al. 2001, Tischer et al. 2009]. Upon such contact-induced catastrophes, proteins that specifically associate with growing microtubule plus ends, so-called +Tips [Akhmanova and Steinmetz 2008], may be transferred to the cortex leaving behind a protein “hot spot” that contributes to the establishment of a cortical protein distribution [Siegrist and Doe 2007]. This has been extensively shown in *S. pombe* [Mata and Nurse 1997, Dodgson et al. 2013], and may occur also in neurons and migrating cells [Horiguchi etal. 2012, Siegrist and Doe 2007].

Although microtubule-mediated protein transfers have been observed *in vivo*, it is not clear what the requirements for successful transfers are. Interestingly, +Tips that are transferred *in vivo* associate with microtubule plus ends by forming complexes with other proteins, including motor proteins. In *S. pombe*, transfer of the protein Tea1, responsible for polarized cell growth, requires the interaction between several +Tips (Figure 1A). These include the EB homologue Mal3, the kinesin-like Tea2, and the CLIP-170 homologue, Tip1, among others [Busch et al. 2004a, Busch et al. 2004b, Martin et al. 2005] (see Figure S1A for more details). Tea1 binds to Tip1-Tea2 complexes by direct interaction with Tip1 [Behrens and Nurse 2002, Feierbach 2004]. Tip1-Tea2 complexes are recruited to the microtubule lattice and transported to the microtubule end with the help of Mal3 [Bieling et al. 2007]. Transfer of Tea1, Tip1, and Tea2, but not Mal3, to the cell cortex occurs by direct interaction between Tea1 and the transmembrane prenylated anchor Mod5 [Behrens and Nurse 2002, Browning 2003, Snaith and Sawin 2003, Busch et al. 2004, Feierbach et al. 2004, Snaith et al. 2005, Meadows et al. 2018]. At the cortex, Tea1 proteins, as well as other cell end factors, have been found in clusters of sizes up to 100 nm [Bicho et al. 2005, Dodgson et al. 2013].

**Figure 1.**
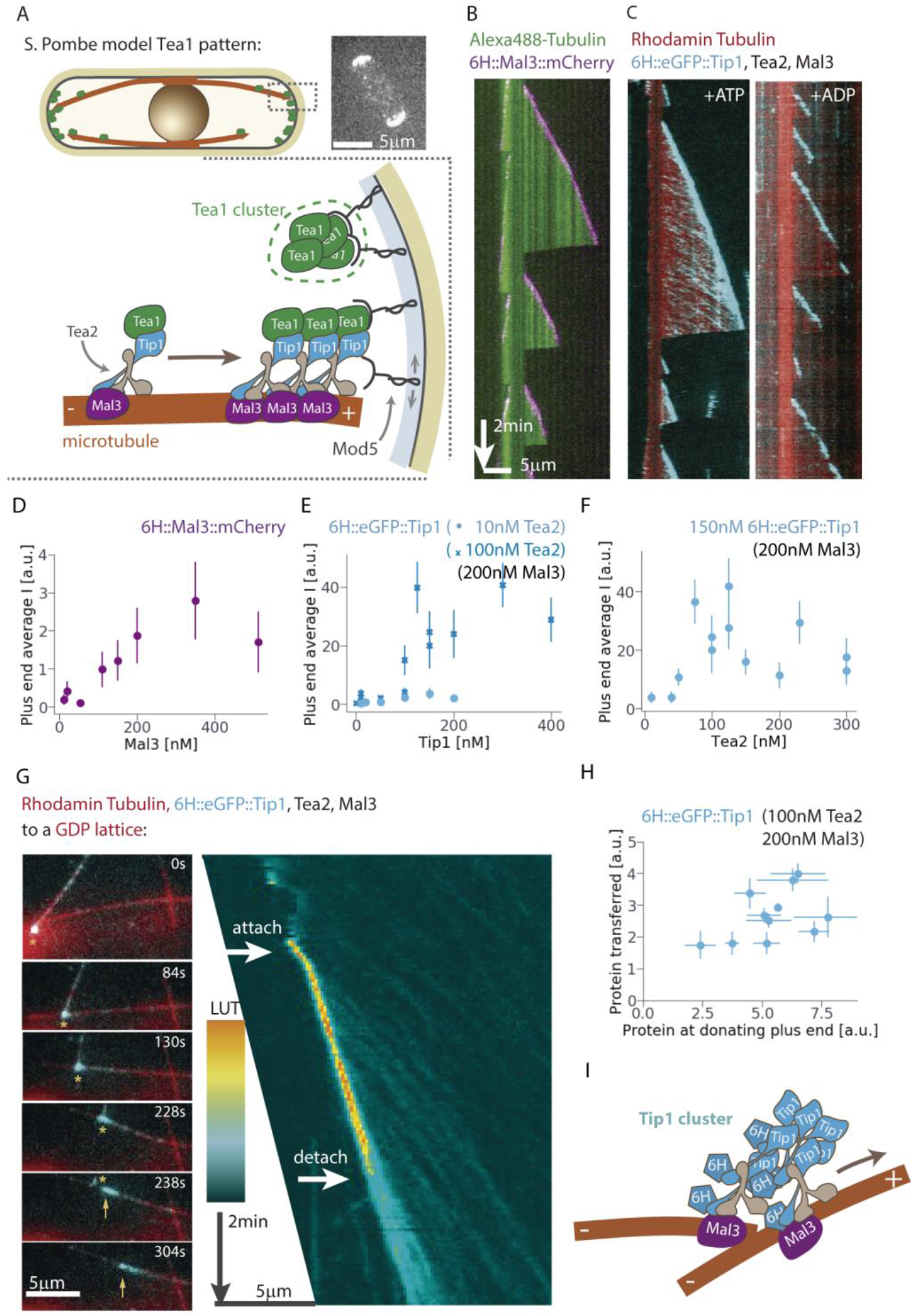
A. Schematic and fluorescent microscopy image of eGFP::Tea1 polarization in *S. Pombe*. Complexes of Tea1, Tip1 and Tea2 found at microtubule ends are transferred to the cell poles via interaction with Mod5. B,C. Kymographs of microtubules grown from GMPCPP stabilized ‘seeds’ in the presence of +Tips *in vitro*. D-F. Average fluorescence intensity at the microtubule plus ends as a function of different protein concentrations. G. Example of a transfer of a 6H::eGFP::Tip1 cluster from a microtubule end to another microtubule lattice and the corresponding kymograph of the fluorescent 6H::eGFP::Tip1 signal along the receiving microtubule. Asterisks and arrows indicate the locations of the microtubule plus end and the transferred cluster of 6H::eGFP::Tip1 respectively. H. Efficiency of multiple transfer events from microtubule plus ends to crossing microtubule lattices. I. Schematic picture of a cluster being transferred from one microtubule to another.

In this study, we sought to establish an *in vitro* assay in which the mechanism of microtubule-mediated protein transfer could be addressed with a minimal number of components. The specific question we asked is whether and how the transfer of +Tips to a biomimetic cortical receptor differs between autonomous plus-end trackers and motor-mediated plus-end complexes. We developed an assay where dynamic microtubules bring His-tagged +Tips in contact with a glass wall functionalized with Ni-NTA (Figure 3A) [Taberner et al. 2014]. We tested under which conditions this microtubule-based delivery of proteins to the wall led to successful transfers. As an autonomous tip tracker, we used His-tagged Mal3, and as a motor-mediated complex we used the Tip1-Tea2-Mal3 system with only Tip1 carrying a His-tag. In both cases, only the His-tagged proteins had affinity for the functionalized walls. We found that His-tagged Tip1 formed clusters at microtubule ends that were efficiently transferred to functionalized walls (as well as other microtubule lattices). In contrast, His-tagged Mal3, which accumulates at microtubule ends but does not appear to form clusters, was not transferred, even when total protein numbers at microtubule ends were comparable. We further observed that in the presence of ADP-bound Tea2, which has reduced microtubule affinity, His-tagged Tip1 cluster formation at microtubule ends was highly reduced and so were transfers to the walls. These observations show that a locally enhanced concentration of proteins at the microtubule end is not enough to produce successful transfers. Instead, clustering at microtubule ends seems necessary. We further found that microtubule catastrophe events, although not necessary, facilitated protein transfers. We finally used the developed *in vitro* assay to show that transfer of protein clusters by microtubule ends is biased to the chamber poles of elongated micro-fabricated chambers, when microtubules spontaneously align. This leads to a mildly polarized protein distribution in a chamber-length-dependent manner.

## Results

### Tip1 forms transferable clusters at microtubule ends

Before confronting growing microtubules with functionalized walls, we screened experimental parameters to find out under which conditions maximum accumulation of His-tagged +Tips at growing microtubule plus ends is achieved. We let microtubules polymerize from GMPCPP stabilized ‘seeds’ bound to a glass slide in the presence of either 6H::Mal3::mCherry or Mal3, Tea2 and 6H::eGFP::Tip1 at different protein concentrations. As shown in earlier studies, 6H::Mal3::mCherry forms comets at growing microtubule plus ends with fluorescent intensities saturating at concentrations around 200nM. (Figure 1B and D) [Bieling et al. 2007, Maurer et al. 2011]. As also previously shown [Bieling et al. 2007], the fluorescent signal of 6H::eGFP::Tip1 (150 nM), in the presence of Mal3 (200 nM) and Tea2 (with ATP), reveals transport traces along GDP microtubule lattices as well as protein accumulation at growing microtubule plus ends (Figure 1C). At low Tea2 concentrations (10 nM), 6H::eGFP::Tip1 accumulation at microtubule ends was comparable to autonomous 6H::Mal3::mCherry accumulation at similar concentrations (Figure 1E). Increasing Tea2 up to 100nM led to a higher accumulation of 6H::eGFP::Tip1 at the plus ends (Figure 1E and F). Note that the Tip1 signal increased in a non-linear way both as a function of Tip1 concentration (Figure 1E) and Tea2 concentration (Figure 1F), and was highly variable at high motor concentrations. Therefore, the number of 6H::eGFP::Tip1 molecules that can accumulate at microtubule ends with the help of active Tea2 is much higher than the number of 6H::Mal3::mCherry molecules that can autonomously track microtubule ends.

In addition we observed, even at low Tea2 concentrations, that 6H::eGFP::Tip1 proteins which accumulated at growing plus ends could be transferred to other crossing microtubule lattices in apparent “clusters” that continued advancing on the accepting lattice (Figures 1G and H). During transfers, the clusters formed physical links between the two microtubules that could last several seconds (Figure 1G). Via this link, a donating microtubule plus end could be stirred along an accepting microtubule (Figure 1G), which suggests that clusters contain multiple motor proteins (Figure 1I). Interestingly, transferred clusters often showed lower speeds than “normal” Tea2-driven Tip1 traces on microtubule lattices (Figure 1G and Supplementary 2). We thus conclude that Tip1 can form transferable clusters at microtubule plus ends, even at low Tea2 concentrations. This motor-mediated cluster formation is likely the reason that at high Tea2 concentrations, Tip1 can accumulate at much higher numbers than saturating numbers of Mal3.

We note that “normal” 6H::eGFP::Tip1 traces were not observed on GMPCPP microtubule lattices (the seeds), for which Mal3 has very low affinity (Figure S1B) [Maurer et al. 2011]. Interestingly, 6H::eGFP::Tip1 transported from growing minus ends detached from the microtubule when reaching the GMPCPP lattice (Figure S1B). This suggests that Mal3 is not only needed for recruitment of Tip1 and Tea2 to the microtubule lattice, but also for transport along the lattice.

### Preparing the biomimetic cortex

To create a biomimetic cortex, we fabricated glass micro-chambers with 300-400 nm high sidewalls, containing a double layer of 50 nm chromium and 50 nm gold on top to create an overhang (Figure 2A, S2AB). The walls of the chambers were functionalized with chelated nickel ions using poly(ethylene glycol) - grafted with copolymer poly(L-lysine), and derivatives of copolymer poly(L-lysine) with a functional three-head nitrilotriacetic (PLL-g-PEG/tris-Ni(II)-NTA) [Bhagawati et al. 2013] (Figure 2B, S2C). The bottom surfaces of the microchambers were coated with either PLL-g-PEG/biotin for subsequent immobilization of biotinylated microtubule GMPCPP ‘seeds’ or PLL-g-PEG for passivation against unspecific absorption of proteins using a previously described method [Taberner et al. 2014].

**Figure 2.**
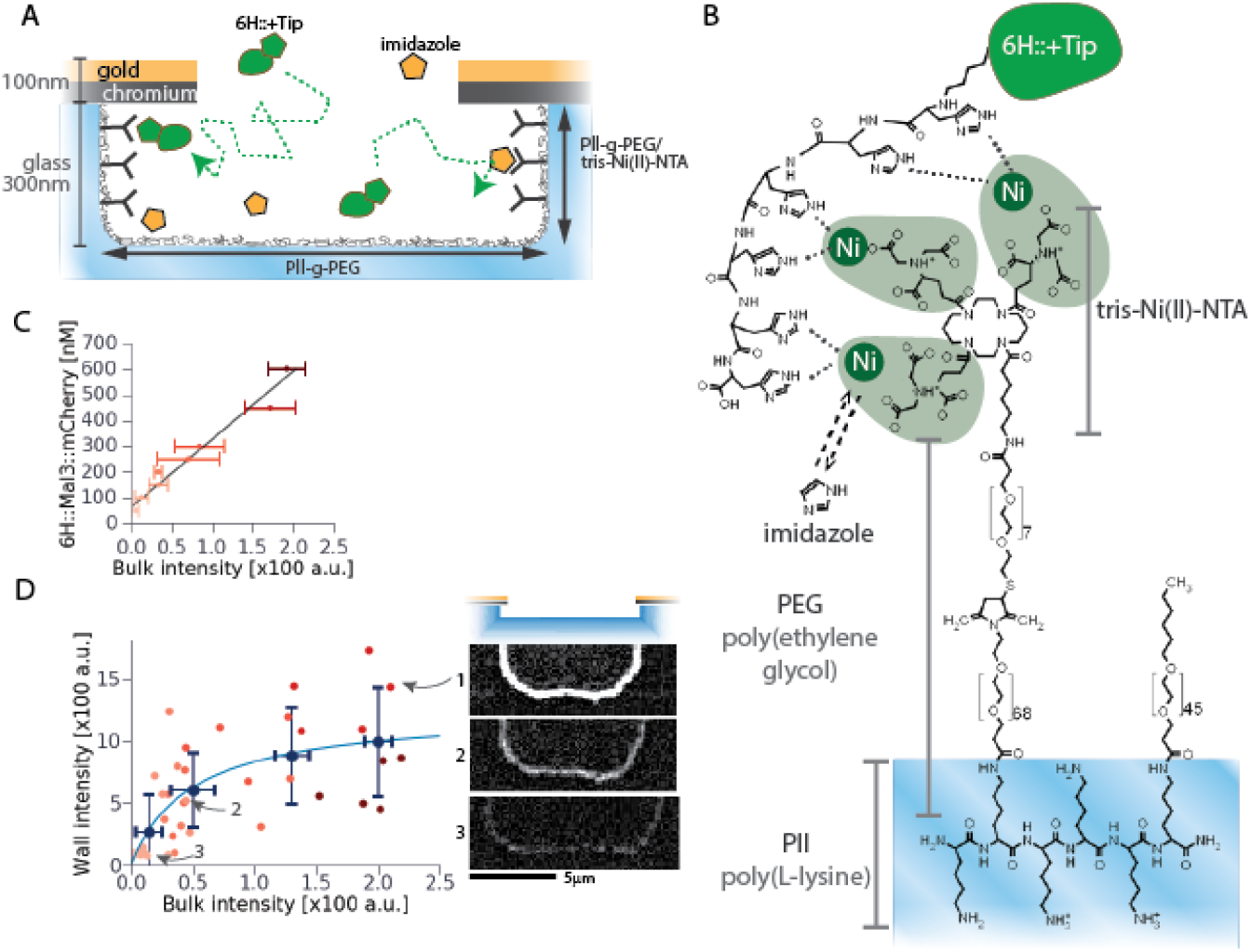
A. Schematic of the microchambers and their surface functionalization. His-tagged proteins can reversibly bind to the walls coated with Pll-g-PEG/tris-Ni(II)-NTA. The presence of imidazole in solution competes for binding. B. Chemical structure and binding of a His-tagged protein to nickel ions chelated by Pll-g-PEG/tris-Ni(II)-NTA. C. and D. Binding of 6H::Mal3::mCherry to micro-chambers with walls coated with Pll-g-PEG/tris-Ni(II)-NTA with 10 mM imidazole. C shows the relation between 6H::Mal3::mCherry protein concentration and the measured bulk fluorescence signal (far from the wall). Error bars correspond to standard deviation. D shows the fluorescence signal at the wall, as a function of the bulk signal. Yellow to red dots correspond to individual measurements (mean value along 1 um of the wall), color coded by protein concentration as in C. Black dots correspond to the binned data by the Freedman-Diaconis rule (bins of 67 a.u.) [Freedman and Diaconis 1981], except for the first bin which is smaller (35 a.u.). Blue line corresponds to a fitted binding curve. The right side shows examples of background subtracted images for individual measurements indicated in D.

Each Tris-Ni(II)-NTA unit contains three chelated nickel(II) ions (Figure 2B). To each of these ions two histidines can reversibly bind via their imidazole functional group. A His-tagged protein, containing six consecutive histidines, can thus bind stoichiometrically to one Tris-Ni(II)-NTA unit [Lata and Piehler 2005]. Addition of imidazole in solution competes with His-tagged proteins for binding sites, therefore it effectively reduces His-tagged protein affinity for Tris-Ni(II)-NTA. To avoid strong direct binding of proteins from solution to the walls, the cortical affinity needs to be slightly lower than that of the proteins for the microtubule ends. We found that at 10 mM imidazole, 6H::eGFP::Tip1 as well as 6H::Mal3::mCherry at 150 nM would exhibit very low binding to the wall, fulfilling this condition. Indeed, measurement of the 6H::Mal3::mCherry binding curve to the wall at 10 mM imidazole yielded a dissociation constant of K_D_ = 190±20 nM, indicative of the concentration at which on average half of the wall binding sites are occupied with 6H::Mal3::mCherry (Figure 2CD). Alternative experiments to measure the K_D_ of 6H::eGFP::Tip1 in individual samples gave similar order of magnitude results (Figure S3).

### Microtubules transfer Tip1 clusters to the biomimetic cortex

We then tested whether microtubule ends could transfer his-tagged Mal3 or his-tagged Tip1 to the functionalized walls. Stabilized microtubule seeds were immobilized on the bottom surface via biotin-streptavidin linkages. Dynamic microtubules were grown in the presence of either 150 nM 6H::Mal3::mCherry or 150 nM 6H::eGFP::Tip1, 10 nM Tea2 and 200 nM Mal3 with an imidazole concentration of 10 mM (Figure 3A). We observed microtubule plus end contacts with the wall and measured the fluorescence intensity at the contact point over time.

**Figure 3.**
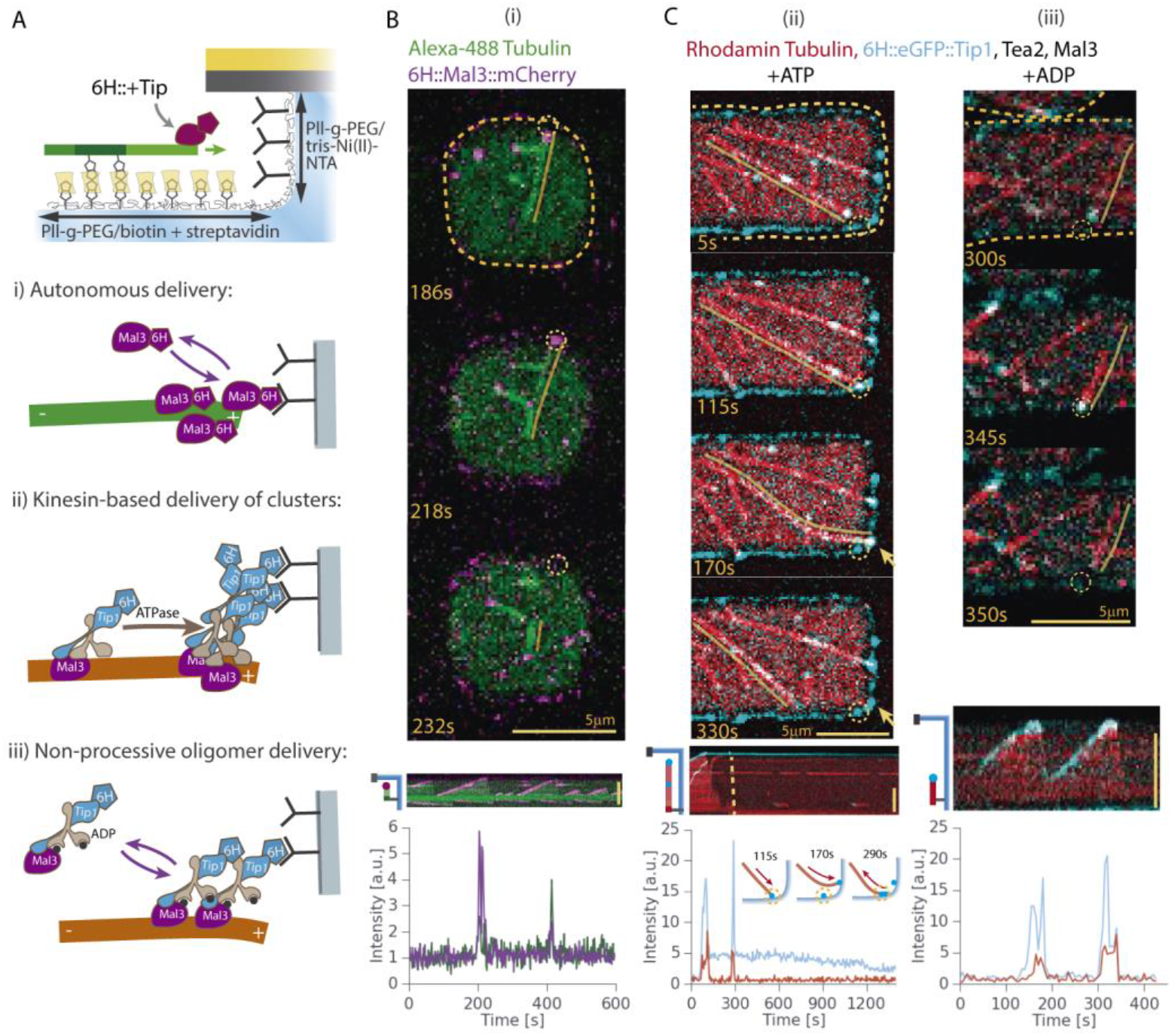
A. Schematic of the experimental setup for testing microtubule-based transfers of different +Tips. B and C. Examples of different deliveries of his-tagged +Tips: 6H::Mal3::mCherry in B, 6H::eGFP::Tip1 in the presence of Tea2 and Mal3 with ATP (left) or ADP (right) in C. Top images show snapshots before, during, and after microtubule tip contact with the wall. The wall location and the initial contact site are indicated by a yellow dashed line and circle respectively. A yellow line has been drawn parallel to the microtubule of interest as a guide to the eye. Arrows point at transfers at subsequent contact sites. The snapshots are accompanied below by a kymograph along the microtubule trajectory of the measured fluorescence intensity over time at the initial contact site. Cartoon insets in the 6H::eGFP::Tip1 case indicate the sequence of events for the microtubule of interest, consisting of an initial growth phase along the wall followed by a shrinking phase.

Figures 3B and Movie 2 show a typical example of a microtubule delivering 6H::Mal3::mCherry to the wall (dotted yellow circle). In these experiments, the protein delivered to the wall did not remain attached after microtubule catastrophe. This can be seen in the kymograph along the microtubule, and also in the fluorescence intensity at the contact point with the wall over time (Figure 3B below). Although the 6H::Mal3::mCherry fluorescence signal at the wall increased upon microtubule contact (at 218 s), it returned to pre-contact values after microtubule catastrophe (at 232 s). Fluorescence intensity values were normalized to the average intensity elsewhere at the wall (where no protein was delivered). In the example shown, the spot on the wall reached by the microtubule contained no pre-bound protein, therefore the base intensity (when no microtubule end is inside the spot) fluctuates around 1 a.u. In the case of microtubules delivering 6H::eGFP::Tip1 to the walls, clear transfers were often observed (Movie 3). Figure 3C left shows an example where the microtubule transferred 6H::eGFP::Tip1 to a spot on the wall (at 115 s, dotted yellow circle), and then continued sliding along the wall. This same microtubule later stopped moving at another spot on the wall (at 170s, yellow arrow), transferred extra protein there, and then underwent catastrophe (dashed yellow line in the kymograph). During the shrinking phase, the microtubule end slid back along the wall, and passed again through the first spot. This contact did not leave extra protein behind at the wall (Figure 3C left at 290 s).

We quantified the success of protein transfers for individual events by plotting the amount of protein (as measured by fluorescence intensity) left behind at the wall after microtubule contact (Figure 4 inset) versus the amount of protein delivered by the microtubule plus end. Protein delivered by the microtubules was determined by the average fluorescence intensity at the wall during microtubule contact. We distinguished between cases where the microtubule left the contact point by undergoing catastrophe (‘MT catastrophe’) or by continuing sliding along the wall (‘no MT catastrophe’). We also distinguished between ‘end-on’ contacts (angle between the microtubule and the wall larger than 15°) and lateral contacts (all other cases).

**Figure 4.**
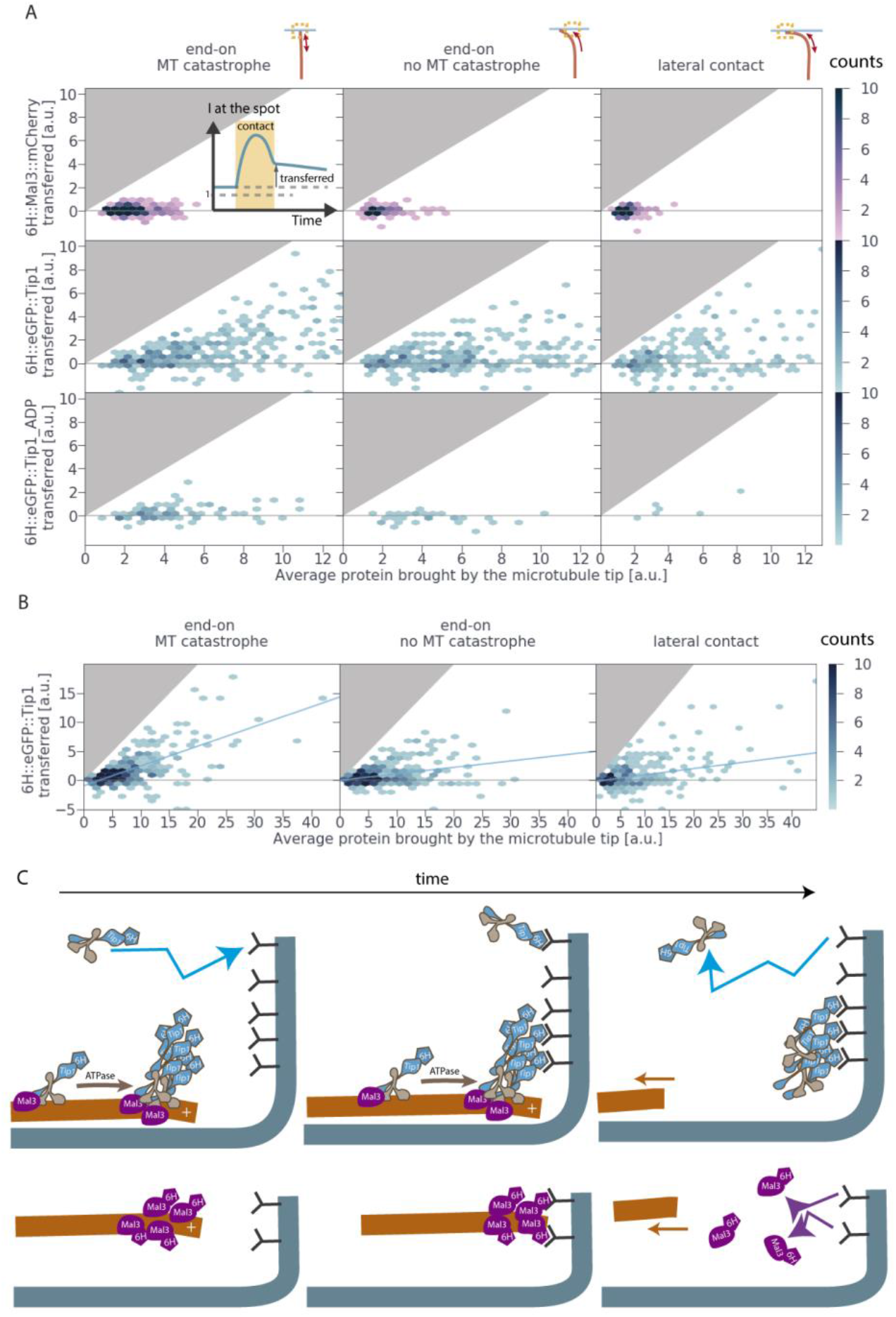
2D histogram plots of efficiency of His-tagged +Tip transfers under different conditions. Detailed explanations can be found in the main text. For the case of 6H::eGFP::Tip1 transfers, linear fits are shown as a guide. A. Close-ups for moderate numbers of proteins at the plus end. B. Full range of 6H::eGFP::Tip1 transfers in the presence of ATP. C. Schematic of motor-mediated clustering of proteins at microtubule ends to facilitate transfers to a biomimetic receptor

Quantification of all plus end contacts with the walls shows that 6H::Mal3::mCherry (Figure 4A first row panels) was not transferred at all, whereas 6H::eGFP::Tip1 was clearly transferred, proportional to the amount of protein delivered by the microtubule plus end (Figure 4A second row panels). His-tagged eGFP::Tip1 transfers exhibited on average 34±1% docking efficiency. Instead, for 6H::Mal3::mCherry the average amount of protein transferred was negligible (^~^0.03 a.u.), regardless of the amount of protein delivered by the microtubule. Many microtubule plus ends delivered more 6H::eGFP::Tip1 protein to the wall than maximally possible for 6H::Mal3::mCherry (Figure 4B) (see also results for freely growing microtubules above). For these high protein amounts, i.e. deliveries above 13 a.u., successful transfers were observed in almost all cases (Figure 4B). Although the example in Figure 3C shows that microtubule catastrophe is not a requirement for successful transfers, many events in which the microtubule left the contact point by sliding along the wall did not lead to successful transfers (Figure 4A and B central panels). As a result, the average efficiency of protein transfer was lower (11±1%) than after catastrophes.

Our observations suggest that Tea2 motor activity promotes the ability of Tip1 to form clusters at microtubule ends, and that cluster formation is a requirement for successful transfers to the functionalized wall. There are however other possible reasons for the difference in behaviour between 6H::Mal3::mCherry and 6H::eGFP::Tip1 proteins. For example, it is possible that aggregates of Tip1 are present in solution which would appear as high-density clusters when transported as cargo by Tea2 motor proteins. In addition, differences in protein size, location of the histidine tag, and exact binding location at the microtubule end [Maurer et al. 2012, Maurer et al. 2014, Zhang et al. 2015] could all lead to differences in the ability of individual proteins to “reach” the functionalized wall while still bound to the microtubule end.

To start with the last possibility, we analysed plus end contacts with the wall in a lateral configuration. In this case, proteins that do not reach the very tip of the microtubule (such as Mal3 under normal circumstances) can still contact the wall. Although lateral contacts led to clear transfers of 6H::eGFP::Tip1, they again did not lead to transfers of 6H::Mal3::mCherry (see Figure 4 A). To rule out the effect of differences in protein size and location of the histidine tag, we next looked for a way to impose Mal3-like microtubule-binding behaviour on Tip1. We removed the microtubule binding and motor activity of Tea2 by substituting ATP by ADP in our 6H::eGFP::Tip1 assays [Cross 2016]. Under these conditions, 6H::eGFP::Tip1 exhibited Mal3-like end tracking behaviour at both microtubule ends and transport traces along the lattice were no longer observed (Figure 1C). In the presence of ADP, Tea2 presumably acts as a passive linker between Mal3 and Tip1 to help recruit Tip1 to growing microtubule ends, since Tip1 does not end track in the presence of Mal3 alone [Bieling et al. 2007 and our own observations]. When we replaced ATP with ADP in 6H::eGFP::Tip1 delivery experiments, clear transfers were rarely observed (Figure 3C, Movie 4, and Figure 4A third row panels). Interestingly, in these assays, 6H::eGFP::Tip1 numbers at microtubule ends were considerably lower than for assays with ATP, indicating that cluster formation had been highly impaired. Since in the presence of ADP the behaviour of Tip1 was similar to that of Mal3, we attribute the failure to transfer to the walls to the inability to form clusters rather than to differences in molecular details. The lack of Tip1 cluster formation in the presence of ADP furthermore argues against the possibility that Tip1 is simply forming aggregates in solution.

### Tip1 clusters stabilize microtubules against catastrophes during wall contact

We observed significantly different dynamics for microtubules encountering the wall in assays with 6H::eGFP::Tip1 containing ATP compared to assays containing ADP, or assays with 6H::Mal3::mCherry. While in the latter two assays 70% of the microtubules underwent catastrophe upon first contact with the wall, this only occurred in 40% of the cases for the 6H::eGFP::Tip1 samples with ATP (Figure 5A, coloured fractions). Microtubules in the remaining (grey) fractions continued sliding along the wall, and eventually underwent catastrophe somewhere else.

**Figure 5.**
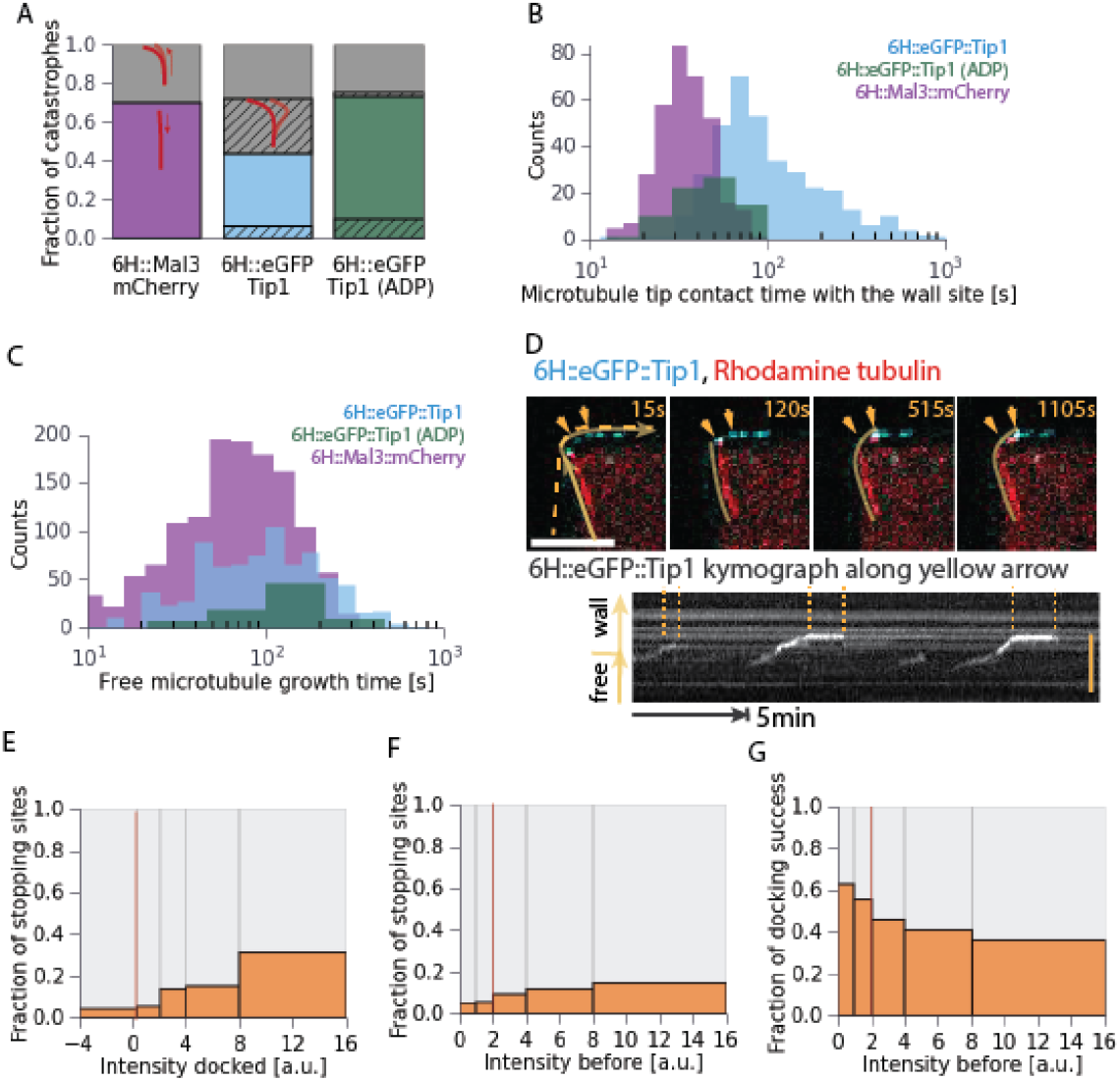
A. Fractions of events where the microtubule undergoes catastrophe at the site of first contact with the wall (colors) versus events where the microtubule first slides along the wall (grey). Dashed areas correspond to fractions of microtubules exhibiting clear buckling due to continued microtubule growth while maintaining contact with a fixed site on the wall. B. Distribution of microtubule end contact times with the wall, with exponential time axis. C. Distribution of microtubule catastrophe times (growth times from seed nucleation to first catastrophe) without wall contact. D. Top panels show snapshots of a microtubule sliding along the wall with its end stopping at two sites (yellow arrows). Dashed yellow line indicates the wall. Continuous yellow line indicates the microtubule. The bottom panel shows a kymograph along the microtubule plus end trajectory (yellow arrows in the first snapshot). This includes a straight segment from the microtubule seed towards the wall, and a curved segment along the wall. Dashed lines indicate the time window in the kymograph where the microtubule end stopped. E. and F. Fractions of pixels along the wall where microtubule plus ends stopped (determined by sliding velocity lower than 0.36 um·min^−1^) as a function of fluorescence intensity change at those pixels (I _docked_ = I _after contact_ - I _before contact_) and fluorescence intensity before the contact respectively. G. G. Fraction of pixels along the wall where microtubules successfully transferred protein as a function of fluorescence intensity at the wall before contact.

We also quantified the fraction of microtubules that showed clear buckling due to prolonged contact with a fixed point on the wall while continuing microtubule growth (Figure 5A, dashed fractions). Only 6H::eGFP::Tip1 samples containing ATP showed significant fractions of microtubule buckling (30% of the events). We previously showed that in the absence of +Tips, microtubules can buckle during growth against a fixed point on the wall [Janson et al. 2003, 2004]. The absence of buckling events in 6H::Mal3::mCherry assays is due to the fact that Mal3 promotes catastrophes in the presence of compressive forces. In the presence of ATP, this effect is counteracted by the presence of Tip1-Tea2 complexes at microtubule contact points with the wall. This is also reflected in the distribution of microtubule-wall contact times (Figure 5B). While in 6H::Mal3::mCherry samples, microtubule ends remained on average 34 s in contact with the wall, microtubules in 6H::eGFP::Tip1 samples with ATP showed an average contact time of 100 s. In the presence of ADP, an intermediate average contact time of 52 s was observed. Instead, quantification of free microtubule catastrophe times (without wall contact) in the same samples showed very little difference between microtubules grown in the presence of Mal3 alone, or with the additional presence of Tip1 and Tea2 (Figure 5C). Therefore, microtubules are specifically stabilized by Tip1 clusters that accumulate at microtubule-wall contact points. The observed Tip1-mediated stabilization of microtubules at the wall is similar to previous *in vivo* observations which showed that ΔTip1 *S. pombe* cells exhibit shorter microtubule plus end contact times with the cell poles than wild type [Tran et al. 2001]. Also, a recent study linked Tea2 with microtubule stabilization in contact with the cell ends [Meadows et al. 2018].

### Tip1 cluster transfer promotes tethering of microtubule ends to the wall

We observed that microtubule ends sliding along the wall often stopped at specific points (Figure 5D). This could be caused by: 1) an irregularity on the wall surface, 2) capturing of the microtubule end by pre-bound 6H::eGFP::Tip1 at the wall, or 3) 6H::eGFP::Tip1 at the microtubule end tethering it to the wall. To distinguish between these possibilities, we measured the fluorescence intensity at the wall before, during, and after microtubule end contact. We then computed the fraction of pixel locations where a microtubule end stopped as a function of the amount of protein deposited (fluorescence intensity after the contact minus intensity before the contact, Figure 5E), and as a function of the fluorescence intensity before the contact (Figure 5F).

The likelihood of microtubule stopping increased with the amount of protein being transferred (Figure 5E). On the other hand, the likelihood of microtubule stopping showed nearly no dependence on the amount of protein pre-bound at the wall (Figure 5F). These data indicate that Tip1 cluster transfer correlates positively with microtubule stopping, suggesting that the transferred clusters can form a physical link between the microtubule end and the wall, similar to the link observed earlier between microtubule ends and crossing microtubules (Fig. 1). Such a link is apparently less likely to be formed by proteins that are pre-bound at the wall than by proteins being delivered by the microtubule. Consistent with this, pre-bound proteins did not appear to provide efficient receptors for clusters delivered by the microtubules. Figure 5G shows that the likelihood of successful docking events in fact decreased with the amount of protein pre-bound at the wall.

6H::eGFP::Tip1 clusters were furthermore occasionally observed to be transferred from the microtubule lattice (far from the microtubule end), and to induce microtubule gliding along the wall (Figure S4). These observations further support the idea that a physical link between the wall and the microtubule can be formed by 6H::eGFP::Tip1 clusters. The observed tethering of microtubule ends by 6H::eGFP::Tip1 clusters at the wall is again very reminiscent to *in vivo* observations. In *S. pombe* cells, microtubule tethering has been suggested to occur via interactions between Tea1 and its cortical receptor, as deletion of Tea1 leads to microtubule curling around the poles [Mata and Nurse 1997].

### Microtubules in elongated microchambers can establish a mildly polarized distribution of Tip1 clusters

Having established that growing microtubule ends can transfer Tip1 clusters to points of microtubule-wall contact, we next tested whether microtubules aligned along the long axis of elongated micro-chambers can establish a polarized Tip1 distribution. In these experiments, microtubules were grown from free GMPCPP seeds inside functionalized rectangular microchambers in the presence of 6H::eGFP::Tip1, Tea2 and Mal3 (and ATP). Micro-chambers were designed with different aspect ratios and sizes ranging from 7 to 50 μm. Microtubule dynamics and 6H::eGFP::Tip1 distributions at the wall were assessed in time-lapse movies over 10-20 minutes (see snapshots in Figure 6A). Since the chambers were not sealed, microtubules could potentially diffuse in and out of the chambers. However, by using methyl cellulose as a crowding agent, microtubules longer than ^~^5 μm remained close to the bottom surface. In addition, the chamber walls were made with an overhang, which prevented microtubules from sliding up the walls.

**Figure 6.**
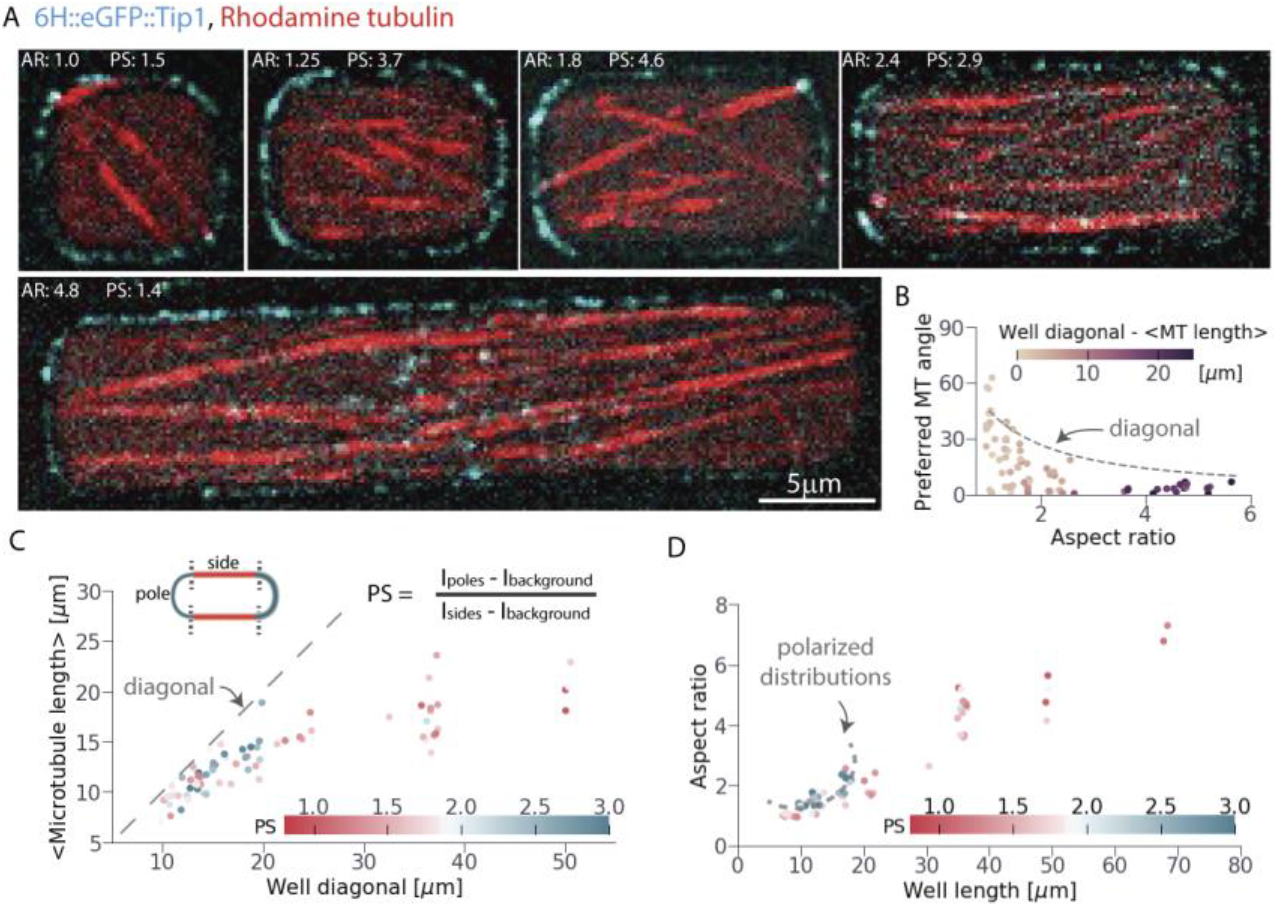
A. Snapshots of dynamic microtubules in microchambers in the presence of 150 nM 6H::eGFP::Tip1, 100 nM Mal3 and 10 nM Tea2. The two laser channels were imaged sequentially with a delay of 400 ms, which led to apparent mismatches between the 6H::eGFP::Tip1 tip signal and the microtubule tip. Aspect ratio (AR) and polarity score (PS) of each chamber is noted. B. Preferred angle of the ensemble of microtubules in each chamber as a function of the aspect ratio of the chamber. C. Mean microtubule length found in each chamber as a function of the diagonal length of the chamber. Color coding shows calculated polarity score (PS). D. Polarity score as a function of the aspect ratio and length of the chamber. Dashed line indicates region in the plot with polarized distributions.

To assess the degree of alignment of the microtubules, we computed the microtubule preferred orientation at each pixel position inside the chambers (Figure S5) and plotted the average preferred angle as a function of aspect ratio for each chamber (Figure 6B). Microtubules in elongated chambers (aspect ratio > 1) preferentially aligned between the longest axis (angle = 0) and the chamber diagonal. Microtubule length distributions depended on the size of the chamber (Figure 6C). For chambers with a diagonal shorter than ^~^20 μm, mean microtubule length scaled with chamber size. However, for longer chambers, microtubule length plateaued at a value of ^~^20 μm. Microtubule lengths in small chambers were likely limited due to catastrophes occurring when the two ends of the microtubule reach the chamber ends. In long chambers, microtubule lengths were instead likely limited by their intrinsic dynamic properties. Microtubules in these long chambers formed two densely packed layers of parallel microtubules (smectic layers [Onsager 1949, Frankel et al. 1988]). Surprisingly, the plus ends of these layers faced each other in the centre of the chamber. It is likely that this configuration resulted from Tip1-Tea2 cluster-mediated sliding between antiparallel microtubules. As a result of this microtubule configuration, microtubule plus end contacts with the ends of the chamber were limited.

To quantify the degree of polarization of the Tip1 distributions on the chamber walls, we used a ‘Polarity Score’ introduced by Sokolowski 2013. This score corresponds to the ratio between the amount of protein found at the poles divided by the amount of protein found at the sides. In our assays protein densities were quantified by fluorescence intensity, and the chamber corners were considered as polar regions (Figure 6C inset). Mild polarity (Polarity Score > 2) could be observed only in chambers of sizes between 10 and 20 μm and aspect ratios between 1.6 and 3 (Figure 6C and D, blue colours). In these chambers, the average microtubule length was close to the chamber size (Figure 6C), implying that microtubule plus ends could reach the chamber poles and undergo catastrophes there. In longer chambers, where microtubules were shorter than the chamber size, a Polarity Score around 1 was observed (red colours). These data suggest that mild polarity resulted from microtubule-based transfers of clusters to the poles of the chambers.

## Discussion

In this paper we set up an *in vitro* assay to establish a minimal mechanism for the microtubule-based transfer of proteins to a cortical receptor. We showed that the local increase of protein density at the microtubule end, as exhibited by EB proteins (6H::Mal3::mCherry in our experiments), is not enough to facilitate transfer to a cortical receptor. Instead, motor-mediated clustering of a cargo protein (6H::eGFP::Tip1 in our experiments) at the microtubule end is sufficient for efficient transfer events. Reduction of protein clustering at the microtubule end by disrupting the transport activity of the motor protein (Tea2) substantially reduced the chances of successful transfer (Figure 4C). Protein transfer appeared to be mediated by simultaneous binding of Tip1 clusters to the microtubule end and the wall, as frequent microtubule end tethering at the contact sites was observed. In addition, although not necessary, microtubule catastrophes increased the amount of protein being transferred to the wall. Finally, we showed that an elongated chamber shape can bias microtubule-based transfers to the poles of chambers with sizes and aspect ratios comparable to *S. pombe*. Below we discuss the relevance of our observations for the *in vivo* establishment of polarized protein distributions.

### The nature and role of motor-mediated clustering at microtubule ends

So far, we did not comment on the nature of the motor-mediated clusters at microtubule ends. Interestingly, the three proteins used, Mal3, Tea2, and Tip1, are all constitutive dimers [Bieling et al. 2007, Sen et al. 2013]. Each of these proteins can directly interact with the other two as well as with the microtubule ([Busch et al 2004a-b, Browning et al. 2005], Figure S1A). Therefore, dynamic protein clusters may be held together by multiple low affinity bonds, templated by the microtubule. In other systems, similar interactions have been found to form liquid-liquid phase separated protein droplets under crowding conditions [Lin et al. 2015, Hernández-Vega et al. 2017, Wang et al. 2018]. In our system, the motor-activity may serve to drive proteins sufficiently close together to enhance such effects. Since depositions at the wall often involve only a fraction of the total amount of protein present at the microtubule end, it is unlikely that single rigid clusters are formed.

What may be the reason that clustering of polarity factors at growing microtubule ends facilitates transfer to cortical receptors? An attractive explanation is that protein clusters that can simultaneously interact with the microtubule end and the receptor, can transfer to the cortex without the need of 3D diffusion. Such a multi-component cluster at the microtubule end may furthermore interact with multiple cortical receptors at the same time. In this way, the cluster at the microtubule end becomes a multivalent ligand which is expected to have a higher collective affinity (avidity) for the cortex than individual polarity factors [Krishnamurthy, et al. 2006]. This would then lead to preferential retention of pre-clustered proteins over individual proteins which arrive by simple 3D diffusion. In this scenario, the role of motor proteins would thus *not* be to transport the proteins all the way towards the receptor, but to facilitate the formation of a protein clusters at microtubule ends that have enhanced affinity for the receptor proteins at the cortex.

*In vivo*, clustering of Tea1 at the cortex was previously shown to be important for cell end deposition [Bicho et al. 2005, Dodgson et al. 2013]. In this case, cluster formation appeared to occur at the plasma membrane as Tea1 gets internalized into a cortical network via interaction with Mod5 [Bicho et al. 2005]. Given our findings, an alternative (or additional) hypothesis may be that Tea1 cluster formation already occurs at the microtubule end, facilitating subsequent transfer to the cell poles.

### Microtubule catastrophes and transfer events

In *S. pombe* cells, Tea1 transfers are typically observed after microtubule catastrophes, but it remains unclear whether there is a strict cause-effect relation. In our *in vitro* setup, his-tagged eGFP::Tip1 transfers were enhanced by microtubule catastrophes, but catastrophes were not strictly required. One can imagine that in a situation where a Tip1 cluster is simultaneously bound to the microtubule end and the functionalized wall, a catastrophe facilitates its transfer by disappearance of the microtubule interaction site. When the microtubule keeps sliding along the wall, binding to the microtubule continues to compete with binding to the wall. In this case, transfer will be less likely, but not impossible.

In our assays, the presence of Tip1 clusters at microtubule ends promoted prolonged microtubule end contacts with the wall. These events included sliding and buckling events and were not always associated with protein transfer. *In vivo*, such stabilization may help reduce microtubule catastrophes at the cortical sides of *S. pombe*, allowing microtubule ends to reach the cell poles [Drummond and Cross 2000, Minc et al. 2000, Tran et al. 2001]. Moreover, the longest microtubule end contacts with the wall were observed in cases of protein transfer. It is therefore possible that simultaneous binding of Tip1 clusters to the wall and the microtubule end leads to further microtubule stabilization. Note that *in vivo*, such long contacts may be prevented by the competing action of destabilizing motors at microtubule ends [Meadows et al. 2018].

### Polarity establishment

In our assays, mildly polarized distributions were established in chambers of length and aspect ratios similar to *S. pombe*. Clearly, the degree of polarization was much less than what is observed *in vivo*. One likely reason for this difference is that microtubule transfers in our *in vitro* experiments were not restricted to the poles of the chamber. In *S. pombe*, the polarized distribution of the receptor Mod5 provides additional positional information. As Mod5 presence at the sides of the cell is very low, transfers there are less probable [Minc et al. 2000, Snaith and Sawin 2003]. We previously showed that an artificial cortical protein that is able to diffuse can form a polarized distribution simply by interactions with microtubule tips in *S. pombe* [Recouvreux et al. 2016]. Therefore, motile receptors, with direct or indirect microtubule binding may facilitate restricting transfers to the poles of the cell. Indeed, receptor redistribution combined with microtubule end capturing, has recently been shown to induce polarity *in silico* [Foteinopoulos and Mulder 2017]. *In vitro*, this combination could be tested by replacing rigid chamber walls with fluid interfaces in water-in-oil droplets with functionalized lipids [Taberner et al. 2015].

Finally, in long micro-chambers (long axis larger than 30 μm), microtubule plus ends showed a clear bias towards the center of the chambers, leading to a failure of cortical polarization. In *S. Pombe* cells, outward microtubule orientation is enforced by Ase1 and Klp2-dependent antiparallel bundle formation [Carazo-Salas and Nurse 2006, Janson et al. 2007]. In the future it will be interesting to test whether such anti-parallel bundles would allow to form polarized distributions in longer micro-chambers as well.

## Materials and methods

### Protein purification

Kinesin Tea2 was expressed in *E. coli* and purified as in [Bieling et al. 2007].

The following proteins were purified as in [Maurer et al. 2011]:

6H::TEV::Mal3::mCherry sequence was obtained by deleting the GFP sequence in 6his::Mal3::GFP (generous gift by Thomas Surrey, [Maurer et al. 2011]) and inserting the mCherry sequence (generous gift from Milena Lanzova and Tom Shimizu) between the *PacI* and *NotI* restrictions sites. To select for correct ligations, we further digested the final constructs with *NdeI* before transformation.

Unlabeled 6H::TEV::Mal3 was obtained as in [Bieling et al. 2007]. The his-tag was removed by cleavage overnight with AcTEV™ protease (Invitrogen, Carlsbad, California, USA) at 4ºC according to the provider’s protocol.

6H::TEV::eGFP::Tip1 was obtained as in [Bieling et al 2007] and the oligo duplex 5’-tccgcgggtgagaatcttcagggcgcc-3’ containing a TEV recognition site was added between the his-tag and the eGFP (recognition sites *SapI* and *NcoI*).

Tubulin was purchased from Cytoskeleton, Inc. (Denver, USA). This includes unlabeled (T240), Rhodamine labelled (TL590M), fluorescent HiLyte 488 (TL488M), and biotinylated (T333P) tubulin from porcine brain.

### Sample preparation

#### Standard flow cell preparation

Glass slides were cleaned by sonication 10 min in water, Hellmanex^®^ (HellmaAnalytics, UK), water, and 70% ethanol sequentially. They were stored in 70% ethanol and sonicated for 10 minutes in water before use.

Coverslips (No.1 24×24 mm x170 μm, Menzel Glässer GmbH, Germany) were cleaned by base piranha (NH_4_OH:H_2_O^2^ in 3:1 at 75°C) for 10 min) and sonicated for 10 min in water.

Flow cells were made by placing two cut 5 mm wide stripes of parafilm on the glass slides separated 5 mm and placing on top the glass slides. Then the parafilm was melted by placing the sample several seconds on a 100°C hot plate and gently pressing a bit on it with tweezers.

**Miro-chamber fabrication** was done as in Taberner et al. 2013 (Figure S2A, see Supplementary information).

#### Micro-chamber functionalization

The microfabricated glass slide was wetted with 0.4 mM Pll-g-PEG/Tris-NTA solution ([Bhagawati et al. 2013], generous gift from Jacob Piehler, Universität Osnabrück, Germany) for 30 min, dried, and exposed to deep UV (185 and 254 nm) with an Ozone Cleaner (Uvotech Systems Corporation, Concord, CA, USA) for 10 min (Figure S2B). Then a flow cell was made with TESA^®^ tape (Tesa SE, Norderstedt, Germany) and the bottom surface was functionalized again by incubating 0.4 mg·ml^−1^ of either PLL(20)-g[3.1]-PEG(2)/PEG(3.4)-biotin(17.5%) or PLL(20)-g[3.1]–PEG(2) (SUSOS AG, Dübendorf, Switzerland) for 30 min. Biotinylated samples were further incubated with 1 mg·ml^−1^ streptavidin for 10 min. All samples were finally passivated with 1.2 mg·ml^−1^ κ-casein.

#### GMPCPP-stabilized microtubule seeds

Microtubule seeds were prepared by a polymerization - depolymerization - polymerization cycle with GMPCPP in MRB80 (80 mM KPipes, 4 mM MgCl^2^, 1 mM ethylene glycol tetraacetic acid (EGTA), pH 6.8). The first polymerization step was performed by incubating a tubulin mix of biotinylated tubulin, fluorescently labelled and non-labelled tubulin in a ratio 18:12:70 (total of 20 μM tubulin) with 1 mM GMPCPP (NU-405 Jena BioScience, Jena, Germany) at 37°C for 30 min. For non-biotinylated seeds the biotinylated tubulin was simply exchanged by non-labelled tubulin. The mix was then airfuged 5 min at 30 psi with an Airfugee^®^ Air-driven ultracentrifuge (Beckman Coulter Inc., Brea, California, USA) and the pellet was re-suspended in 80% of the initial volume and left on ice for 20 min to depolymerize the microtubules. Then, 1 mM GMPCPP was added to the mix and left for 30 min at 37°C to polymerize again. The seeds were then airfuged again, re-suspended in MRB80 with 10% glycerol and stored at 80°C. Thawed seeds were kept at room temperature for up to a week.

#### Tip tracking assays

Flow cells were functionalized with 0.2mg·ml^−1^ PLL(20)-g[3.1]–PEG(2)/-PEG(3.4)–biotin(17.5%) (SUSOS AG), with a subsequent incubation of 0.1 mg·ml^−1^ streptavidin, and further passivation with 1.2 mg·ml^−1^ κ-casein. Biotinylated GMPCPP ‘seeds’ where incubated for 5 minutes. Then, dynamic microtubules where grown from a mix containing 14.25 μM tubulin, 0.75 μM rhodamine tubulin, 50 mM KCl, 0.6 mg·ml^−1^ κ-casein, 0.4 mg·ml^−1^ glucose oxidase, 50 mM catalase, 0.1% methylcellulose, 1 mM GTP, and 2 mM ATP in MRB80 and the variable stated concentrations of 6H::eGFP::Tip1, Mal3, and Tea2.

Tip tracking assays were recorded with a TIRF microscope (Nikon Eclipse Ti-E inverted microscope) equipped with an Apo TIRF 100×1.45 N.A. oil objective, a Perfect Focus System (PFS), a motorized TIRF illuminator (Roper Scientific, Tucson, AZ, USA) and a QuantEM:512SC EMCCD camera (Photometrics, Roper Scientific). Images were taken at 300 ms exposure for excitation lasers 491 nm (40 mW) Calypso (Cobolt, Solna, Sweden) diode-pumped solid-state laser, 561 nm (50 mW) Jive (Cobolt), and 641 nm 28 mW Melles Griot laser (CVI Laser Optics & Melles Griot, Didam, Netherlands) every 2 to 5 s with the same power for all samples. The temperature was kept at 26°C with a homemade objective heater device.

Protein transfer assays were performed in microfabricated glass microchambers with a 150 nM 6H::eGFP::Tip1, 10 nM Tea2 and 200 nM Mal3 and no imidazole. Transfers were recorded as the tip tracking assays with TIRF with a frame rate of 2.0 s.

#### Free binding assay

Functionalized and passivated samples, were incubated with the stated protein concentration of either 6H::Mal3::mCherry or 6H::eGFP::Tip1 for 50 min in a MRB80 solution containing 0.1 % methyl cellulose, 0.4 mg·ml^−1^ glucose oxidase and catalase, 50 mM glucose, 0.6 mg·ml^−1^ κ-casein, 50 mM KCl and 10 nM imidazole. For the equilibrium binding curve fluorescence microscopy images were taken 50 minutes after protein incubation. For the unbinding measurement, the flow cell was flashed with MRB80 buffer with 50 mM KCl and 10 mM imidazole every 5 minutes. For all measurements fluorescence microscopy snapshots were taken at separate locations along the sample at the z position of highest protein fluorescence intensity at the wall. Imaging was done with **spinning disc fluorescence microscopy** using a IX81F-ZDC2 microscope (Olympus, Tokyo, Japan) with a spinning disc confocal head CSU-X1 (Yokogawa, Yokogawa, Japan), 100X oil immersion objectives, and EmCCD camera iXon3 (Oxford instruments, Oxford, UK). Excitation lasers 488 and 561 nm (Oxford instruments) were used. AOTF laser intensity of 22, exposure of 300 ms, and EM Gain 115 were used.

#### +Tip deposition assays

Each functionalized glass slide was used for three flow cells and imidazole concentration was adjusted depending on the level of free his-tagged protein binding to the walls from solution in the first two flow cells. Both, Mal3 and Tip1 assays were prepared in an MRB80 solution containing 14.25 μM tubulin, 0.75 μM labelled tubulin, 0.1 % methylcellulose, 0.4 mg·ml^−1^ glucose oxidase, 50 mM glucose, 0.6 mg·ml^−1^ κ-casein, 50mM KCL, and 2 mM of ATP or ADP when needed. To reach nearly equilibrium state of free protein binding from solution samples were let at imaging temperature for more than half an hour. Measurements of Mal3 and Tip1 depositions were done with the spinning disc microscope mentioned above. Z-stacks of 2-3 focal planes separated 300 nm were taken every 2 s for Mal3 assays and 5 s for Tip1 assays for 25 min.

#### Polarity assessment assay

After functionalization of the walls with Pll-g-PEG/Tris-Ni(II)-NTA (0.2 mg·ml^−1^), the bottom surface was passivated with Pll-g-PEG (0.2 mg·ml^−1^ for 15 min), and κ-casein (1.2 mg·ml^−1^ for 10 min). Then microtubule ‘seeds’ were incubated in an MRB80 solution containing 0.1 % methylcellulose and let sediment for 5 min. The 6H::eGFP::Tip1 deposition assay was prepared by incubating 150 nM 6H::eGFP::Tip1, 100 nM Mal3 and 10 nM Tea2 in a MRB80 solution containing 14.25 μM tubulin, 0.75 μM Rhodamine tubulin, 1 mM GTP, 2 mM ATP, 0.1 % methylcellulose, 0.4 mg·ml^−1^ glucose oxidase and catalase, 50 mM glucose, 0.6 mg·ml^−1^ κ-casein, 50 mM KCl and 10 mM imidazole. Z-stacks of 2-3 focal planes separated 300 nm were taken every 5 s.

## Acknowledgements

We thank Changjiang You and Jacob Piehler for providing the Tris-NTA moieties. We thank the clean room facility of AMOLF (Gijs Vollenbroek, Dimmitry Lamers, Andries Lof, and Hans Zeijlemaker) for support and advice with microfabrication. We thank Vanda Sunderlíková, Magdalena Preciado-Lopez, Sophie Roth, Cristina Manatschal, and Michel Steinmetz for help with protein purification. We thank Liedewij Laan, Pierre Recouvreux, Louis Reese, Maurits Kok, Renu Maan and Vladimir Volkov for discussions, and Louis Reese for critical reading of the manuscript. This work is part of FOM Programme nr. 110, which is financially supported by the “Nederlandse Organisatie voor Wetenschappelijk Onderzoek (NWO).”

---

**Movie 1.**

Transfer of 6H::eGFP::Tip1 (cyan) from a microtubule tip to another microtubule lattice (Rhodamine tubulin in red) and its seed (HiLyte-488 tubulin in cyan) in the presence of unlabeled Tea2 and Mal3.

**Movie 2.**

Microtubule-based deliveries (green) of 6H::Mal3::mCherry (magenta) to Tris-Ni(II)-NTA functionalized walls. No protein is transferred to the walls.

**Movie 3.**

Microtubule-based deliveries (red) of 6H::eGFP::Tip1 (cyan) to Tris-Ni(II)-NTA functionalized walls in the presence of Mal3 and Tea2 and ATP. Multiple successful transfers to the wall can be observed.

**Movie 4.**

Microtubule-based deliveries (red) of 6H::eGFP::Tip1 (cyan) to Tris-Ni(II)-NTA functionalized wall in the presence of Mal3, Tea2, and ADP. No protein is transferred to the walls.

