## Supplementary Figures for "Motor-mediated clustering at microtubule plus ends facilitates protein transfer to a bio-mimetic cortex"

**Figure S1.**

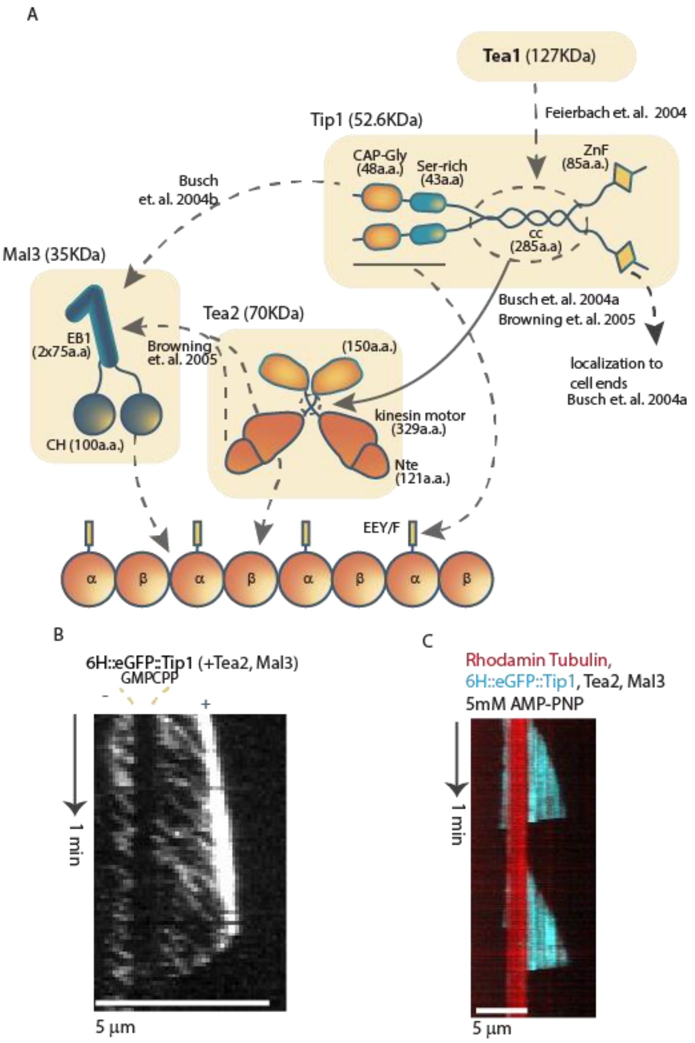


A. Schematic of protein domains and their reported interactions. Dark lines represent unstructured or coiled-coil domains. Dashed and continuous arrows show weak and strong interactions between domains respectively. The exact configuration of the three protein complex on the microtubule is unknown.

B. Close up of the 6H::eGFP::Tip1 signal for 100 nM 6H::eGFP::Tip1, 10 nM Tea2, and 100 nM Mal3 on a kymograph showing no walks on the GMPCPP ‘seed’ and no 6H::eGFP::Tip1 accumulation on the minus side of the ‘seed’.

C. Kymograph of a microtubule with 150 nM 6H::eGFP::Tip1, 10 nM Tea2, and 100 nM Mal3 with 5 mM AMP-PNP instead of ATP.

**Figure S2.**

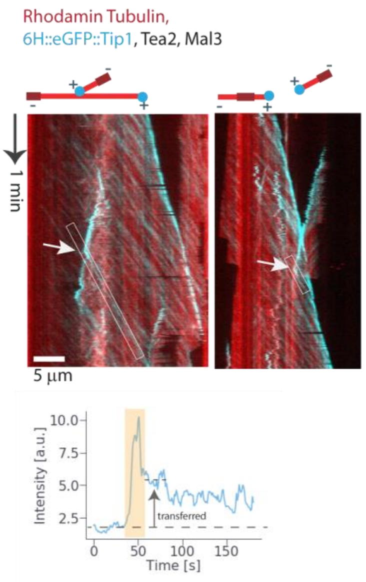


Examples of 6H::eGFP::Tip1 transfers from one microtubule plus end to another microtubule lattice.

Corresponding measurement of transferred protein (left case).

**Figure S3**

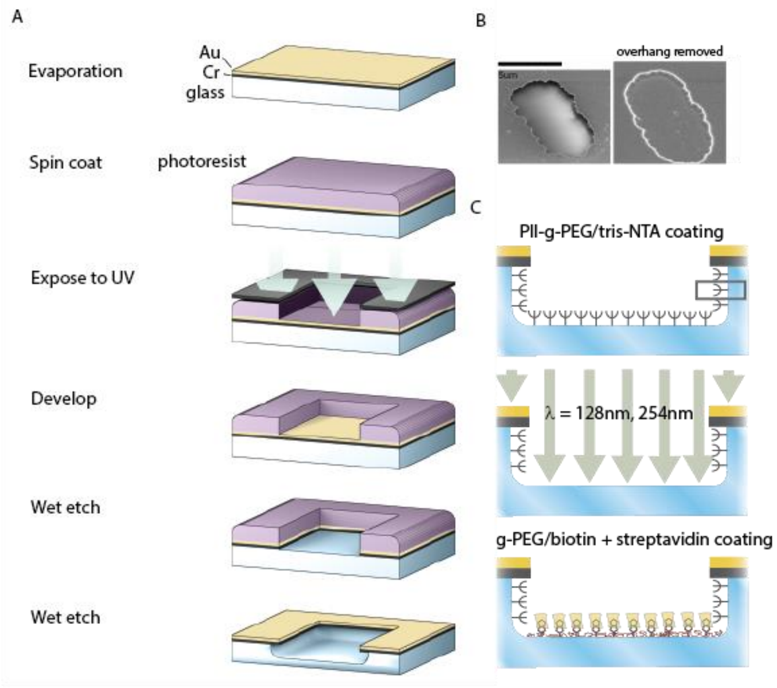


A. Micro-fabrication steps from top to bottom.

B. Surface electron microscopy images of a micro-chamber of a size similar to *S. Pombe*. Right image shows a chamber were the gold and chromium overhangs have been etched to show the glass walls.

C. Selective functionalization of glass microchambers with an overhang. (Top) Coating of all glass surfaces with Pll-g-PEG/Tris-NTA. (Middle) Photo-cleavage of moieties at the bottom surfaces. (Bottom) Coating of the bottom surface with Pll-g-PEG/Biotin for future absorption of streptavidin, followed by biotinylated microtubule 'seeds'.

**Figure S4.**

A

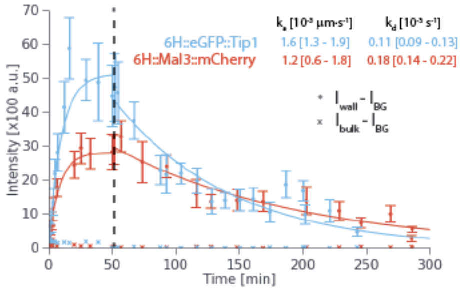


A. Binding kinetics of 1.5 uM 6H::eGFP::Tip1, cyan, or 6H::Mal3::mCherry, red, to Tris-Ni(II)-NTA coated walls at 10 mM imidazole. From time 50 min onwards, unbound protein is removed by flushing 10 mM imidazole every 5 min. Kinetic parameters were obtained by least-square fitting of the binding curve $\Gamma(\mathrm{t}) = \Gamma_{\mathrm{eq}}\left(1 - \mathrm{e}^{-(\mathrm{k_a c} + \mathrm{k_d})\mathrm{t}}\right)$ and unbinding curve $\Gamma(t) = \Gamma_{eq} e^{-k_d t}$. Values in brackets indicate the 95% confidence intervals. Fitting and confidence intervals were obtained with *lsqcurvefit* and *nlparci* of **Matlab R2013b** respectively.

**Figure S5.**

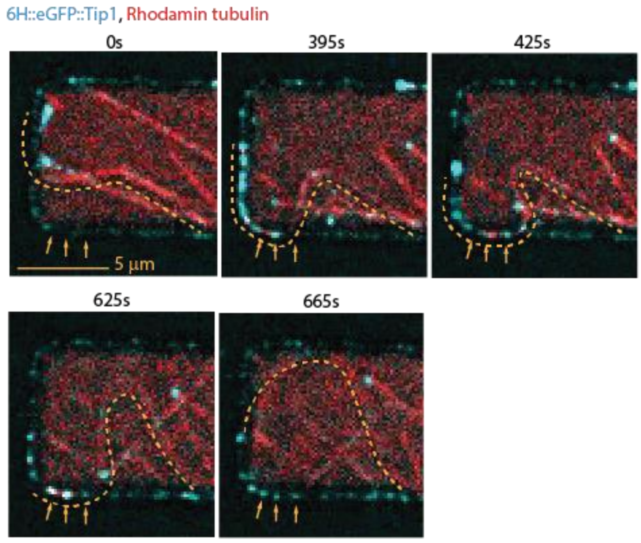


Example of 6H::eGFP::Tip1 transfers (yellow arrows) from the GDP microtubule lattice. Transfers were followed by microtubule gliding at the wall, indicative of a physical link between motor-containing Tip1 clusters at the wall and the microtubule. Yellow dotted lines indicate the shape of the microtubule of interest.

**Figure S6.**

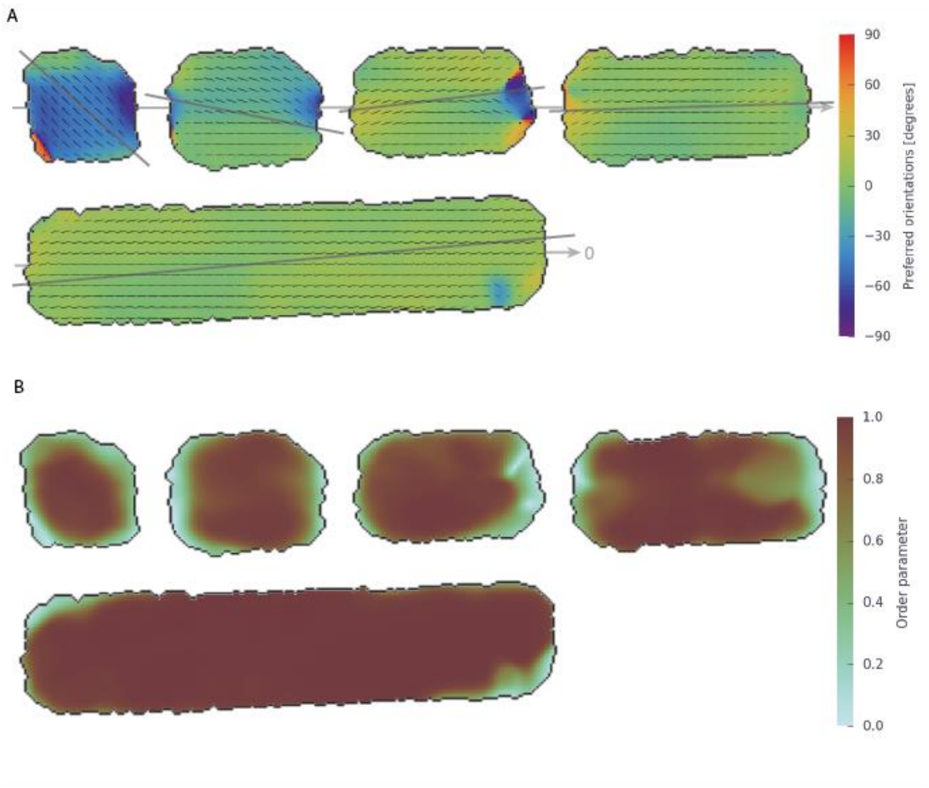


A. Preferred microtubule orientations over the whole 5-10 min movie at each pixel position relative to the long axis of the chamber. The length of the lines is proportional to the order parameter. Grey lines indicate the overall preferred orientation of the ensemble of microtubule in each chamber.

B. Order parameter at each pixel position indicating how probable the preferred angle is.

### Supplementary material:

#### Miro-chamber fabrication:

First, consecutive 50 nm chromium and 50 nm gold layers were evaporated on base piranha cleaned coverslips at a rate of 0.06 $nm \cdot s^{-1}$ in a home-made evaporation chamber at pressure below $10^{-6}$ mbar. Then, a thin layer of HMDS primer (Sigma-Aldrich, USA) was spin coated with a Delta 80 GYSET® Spin coater with a closed lid (Süss MicroTec, Germany) at 4000 rpm for 32 s and baked 1 min at 150˚C. A thin layer (300 nm) of Shipley Microposit® S1805 G2 positive UV-resist (Microresist, Germany) was spin-coated at 500 rpm for 8 s followed by 4000 rpm for 32 s. Then, let dry for 10 s and baked at 115˚C for 60 s. The photo-resist was partially exposed to a dose of 25 $mJ \cdot cm^2$ 365 nm UV light using a Süss MJB3 mask aligner with a binary chromium/soda lime mask on top (Delta mask, The Netherlands). The UV-exposed pattern was then developed for 1 min in Microposit® MF®-319 developer (Microresist).

The gold and chromium layers were subsequently etched with standard gold and chromium etchant respectively (Sigma-Aldrich). The glass chambers were etched isotopically with 40% KOH solution at 80˚C. The sample was cleaned again with base piranha before functionalization.

#### Image analysis:

**Free binding:** Bulk and background intensities ($I_{Bulk}$ and $I_{BG}$) were measured from an area of 1 $\mu m^2$ on the middle of the chambers and on the chromium covered regions respectively in raw images using **ImageJ**. All measurements were performed within the brightest area on the field of view. $I_{Bulk}$ - $I_{BG}$ was then fitted by a linear relation to the protein incubation concentration with the package *numpy.polyfit* of **Python2.7**. Standard deviation errors were obtained as the square root of the diagonal of the covariance matrix obtained from the fitting. Wall intensities were measured from background subtracted images to reduce the effect of non-uniform illumination. Background was estimated by morphological opening of each picture with a 2D disk structuring element of radius 3 (*mahotas.morph.open* and *pymorph.sedisk* packages of **Python 2.7**). Measurements were obtained as the average and std values on 1 μm ROI distance along the brightest wall of the fluorescence image (using **ImageJ**). Background intensities were measured again in background subtracted images. Dissociation constant was obtained by fitting $I^*_{Wall} - I^*_{BG} = \frac{I_{Bulk} - I_{BG}}{I_{Bulk} - I_{BG} + K_D}$ with the package *scipy.optimize.curve_fit* of **Python2.7**. The symbol * denotes that those measurements were performed in background subtracted images. Standard deviation errors were obtained by the square root of the diagonal of the covariance matrix obtain from the fitting.

**+Tip depositions:** For Mal3 assays, a Fourier transform image correction was done to remove periodic speckles that occurred due residual light going through the spinning disk setup. The correction consisted on removing bright spots appearing in the Fourier transform image (**ImageJ** macro with the functions *FFT*, *make rectangle*, *cut*, and *Inverse FFT*). All images were background subtracted at each frame as before and all planes were projected by maximum pixel intensity. All the following image analysis was performed in custom programs on **Python 2.7** in a semi-automated way aided by *PyQt4* graphical user interfaces (GUI). Spots on the wall with protein docked where semi-automatically detected by thresholding of the chromium sheltered region of each micro-chamber. The regions of the wall that were not identified as spots with protein were used to determine the average fluorescence of a site without immobilized protein, which was used for normalization. Spots were automatically distinguished by the contact or not of a microtubule tip at any time during the time lapse movie. Spots with a clear microtubule arrival and departure were manually selected for measurement of the fluorescence intensity over time. To correct for possible drifting, the intensity of the deposited protein was measured in a disk area of 6 pixels radius (*pymorph.sedisk*) centred by a 2D Gaussian fitting of the signal at each frame (*scipy.optimize.curve_fit*) on a 4x interpolated image (*scipy.ndimage.interpolation.zoom*). The exact instants of microtubule arrival and departure were determined manually by looking at the movies. The information of each microtubule contact was stored as a *Python dictionary*. This included the date and movie recording, spot location over time, the times of arrival and departure of a microtubule, the protein fluorescence signal over time, and the conditions in which the microtubule contacted the wall; i.e. whether it was a lateral or a tip contact, if the microtubule underwent catastrophe of not, and the count of times a microtubule had visited that same spot before in the movie. Information of all the microtubule contact events was then organized in *Panda*'s *DataFrame* structures and plot selectively according to its specifications. The average fluorescence before and after microtubule contact were calculated from the 4 and 2 time measurements right before and after microtubule contact (20 s and 10 s for 6H::eGFP::Tip1 deposition experiments and 8 s and 4 s for the 6H::Mal3::mCherry experiments). The comet intensity was measured by Gaussian fitting of the fluorescence signal during microtubule contact (*scipy.optimize.curve_fit*). The linear fitting to obtain the deposition efficiency was obtained by using the package *scipy.stats.linregress*.

**Kymograph analysis** was performed by drawing in **ImageJ** 3 pixel wide splines on the path performed by the microtubule end along the wall. Then, the functions *Straighten, Reslice* and *Z Project* were used. Microtubule end position at the kymographs was then traced manually on the obtained kymograph. Intensities before, during and after microtubule contact where determined automatically with a custom **Python2.7** program. Intensity of microtubule contact corresponded to the average of 5-time pixels on the kymographs centred on the manually traced line. Intensities before and after corresponded to the previous and posterior 4-time pixels. The background was obtained from a close-by region without pre-docked protein.

**Polarity assessment:** Obtained z-stacks were background subtracted and 2D projected by maximum pixel intensity as explained before. Part of the image analysis performed was based on a previous method from Alvarado et al. 2014. Microtubule orientations were obtained by employing the plugin *OrientationJ* from **ImageJ** (http://bigwww.epfl.ch/demo/orientation/\#analysis, [Fonck et al. 2009, Rezakhaniha et al. 2011]). To avoid including the wall orientation in the measurement, chambers were segmented and the background outside signal was replaced by a Gaussian noise with the same mean and std as the signal inside the chambers (Figure S6). Chamber segmentation was performed by semi-automated threshold using the packages *mahotas* and **scipy.ndimage.label** from **Python2.7**.

The orientation of the long axis of the chambers was obtained by *skimage.measure.orientation* on the labelled longest chamber of the field of view and assumed parallel for all the other chambers. Then, local anisotropies where measured with *OrientationJ* using a Gaussian window with radius $\sigma$ = 3 px for each frame. The program gives a matrix with the orientations at each pixel position and the coherency matrix, a measure of the degree of co-alignment with the neighbouring pixels.

The preferred orientation of microtubules at each pixel position over a whole movie was obtained by building the weighted second-order tensor order parameter $S_2$ [Hess et al. 1980] at each pixel location (*px*)

$$S_{2,px} = \begin{pmatrix} \frac{\sum_{i=1}^{n} \omega_{i,px} \cos 2\theta_{i,px}}{\sum_{i=1}^{n} \omega_{i,px}} & \frac{\sum_{i=1}^{n} \omega_{i,px} \sin 2\theta_{i,px}}{\sum_{i=1}^{n} \omega_{i,px}} \\ \frac{\sum_{i=1}^{n} \omega_{i,px} \sin 2\theta_{i,px}}{\sum_{i=1}^{n} \omega_{i,px}} & -\frac{\sum_{i=1}^{n} \omega_{i,px} \cos 2\theta_{i,px}}{\sum_{i=1}^{n} \omega_{i,px}} \end{pmatrix}$$

where $\theta_{i,px}$ denotes the orientation at that pixel at time frame i, and $\omega_{i,px}$ denotes the respective coherency. The eigenvalues of this tensor give the order parameter

$$\lambda_{\pm} = \pm OP_{px} = \pm \sqrt{\left(\frac{\sum_{i=1}^{n} \omega_{i,px} cos\ 2\theta_{i,px}}{\sum_{i=1}^{n} \omega_{i,px}}\right)^2 + \left(\frac{\sum_{i=1}^{n} \omega_{i,px} \sin 2\theta_{i,px}}{\sum_{i=1}^{n} \omega_{i,px}}\right)^2}.$$

The preferred orientation is pointed by the eigenvector $\overrightarrow{\lambda_+} = \lambda_{+,x}\vec{x} + \lambda_{+,y}\vec{y}$ corresponding to the eigenvalue $\lambda_+$. The preferred angle, $\langle\theta\rangle_{px}$, is obtained by

$$\langle\theta\rangle_{px} = \arctan\left(\frac{\lambda_{+,x}}{\lambda_{+,y}}\right)_{px}.$$

The eigenvalues and eigenvectors of $S_{2,px}$ were obtained using the function *numpy.linalg.eig* from **Python2.7**. The relative preferred angles of the microtubules at each pixel location were obtained by subtracting the previously obtained orientation of the long axis to $\langle\theta\rangle_{px}$.

The preferred orientation of the ensemble of microtubules inside each chamber was determined by calculating now the following tensor

$$S_{2,well} = \begin{pmatrix} \frac{\sum_{j=1}^{m} OP_j \cos 2\langle\theta\rangle_j}{\sum_{j=1}^{m} OP_j} & \frac{\sum_{j=1}^{m} OP_j \sin 2\langle\theta\rangle_j}{\sum_{j=1}^{m} OP_j} \\ \frac{\sum_{j=1}^{m} OP_j \sin 2\langle\theta\rangle_j}{\sum_{j=1}^{m} OP_j} & -\frac{\sum_{j=1}^{m} OP_j \cos 2\langle\theta\rangle_j}{\sum_{j=1}^{m} OP_j} \end{pmatrix}$$

where j corresponds to each pixel location of a chamber compost of $m$ pixels. The preferred orientation of the chamber is obtained from linearisation of the matrix as before.

**Microtubule lengths** were measured by direct tracing with **ImageJ** in some frames.

**The polarity score** was obtained by first computing the fluorescence intensity profile of protein at the wall as a function of the distance from the pole. For this, the signal 1 µm away from the walls was masked using a threshold detection (Figure S7, top). Then, we obtained kymographs at each $y$ position parallel to the long axis (arrow in Figure S7 top) using the functions *Straighten* and *Reslice* from **ImageJ**. We computed the average intensity (Figure S7 bottom) at each $x$ position over time. We defined the background as 40 a.u, slightly lower than any measured intensity value, and computed the average intensity at the poles and the sides, delimited by the first and last 10 pixels. The polarity score was calculated by applying $\text{Polarity score} = \frac{I_{poles} - I_{background}}{I_{sides} - I_{background}}$.
